# Computer-assisted dissection of the DNA mismatch recognition mechanism of the MutS protein

**DOI:** 10.64898/2025.12.16.694542

**Authors:** Wenyan Jiang, Yiyang Ding, Yuxuan Li, Haojie Sun, Xiao Liang, Quanshun Li

## Abstract

By sensing local conformational strains in DNA, *Taq*MutS utilizes a push-pull mechanism *via* the synergy of residues S57 and D472 to modulate π-stacking interactions between the key recognition residue F39 and DNA, thereby achieving specific mismatch-site recognition. Meanwhile, residues K61 and R473 fine-tune non-specific binding of *Taq*MutS through electrostatic synergy.

---

Owing to its ultrahigh storage density, favorable chemical stability and low energy consumption for storage, DNA has emerged as a next-generation storage medium that transcends the physical limits of conventional storage technologies.^1-4^ However, side reactions occurring during DNA synthesis can induce an error rate as high as 1-3 errors per kilobase, which not only drastically impairs synthesis efficiency but also drives up production costs.^5,6^ Furthermore, DNA synthesis errors increase exponentially with fragment length, resulting in a low fidelity rate of long-oligonucleotide pools.^7,8^ Therefore, the adoption of error-correction technologies to reduce synthesis error rates represents an effective strategy to improve DNA synthesis quality and cut production costs.^9^ Currently, molecular weight-based screening methods such as high-performance liquid chromatography and polyacrylamide gel electrophoresis can efficiently eliminate length-mismatched erroneous sequences. Nevertheless, these approaches are labor-intensive and ineffective against minor errors such as single-base deletions, insertions or substitutions.^10,11^ In contrast, the mismatch-binding protein MutS-mediated error-correction method enables the isolation of multiple error types from DNA synthesis libraries using a single protein, thus exhibiting enormous application potential in high-fidelity DNA synthesis.^12^

Currently, two major MutS proteins are employed for error correction in DNA synthesis, namely *Taq*MutS derived from *Thermus aquaticus* and *Eco*MutS from *Escherichia coli*.^13-16^ Among them, *Eco*MutS exhibits superior specific mismatch recognition capability over *Taq*MutS.^15^ However, *Eco*MutS shows poor thermostability, as its error correction efficiency drops to less than 90% of the initial activity after 21 days.^17,18^ In contrast, *Taq*MutS possesses higher stability and can retain its original activity over a broad pH and temperature range. Nevertheless, *Taq*MutS displays pronounced non-specific recognition during DNA error correction, which leads to the erroneous elimination of perfectly paired DNA strands misidentified as mismatched sequences.^9^ Therefore, targeted reduction of its non-specific recognition and enhancement of specific binding efficiency can improve the precision of the *Taq*MutS error correction system, providing a more efficient tool enzyme for error correction in DNA synthesis.

*Eco*MutS and *Taq*MutS were expressed in *Escherichia coli* BL21(DE3) and purified by affinity chromatography, followed by assessment of purity using SDS-PAGE. A single, sharp band with molecular weights of 98 kDa (*Eco*MutS) and 95 kDa (*Taq*MutS), respectively, was observed in the 150 mM imidazole elution fractions (**Fig S1** and **S2**), consistent with their theoretical molecular masses. These results indicated that *Eco*MutS and *Taq*MutS were obtained with high purity, providing a solid basis for subsequent functional studies. To systematically compare the mismatch recognition characteristics of *Eco*MutS and *Taq*MutS towards different DNA mismatch types, six typical error types encountered in DNA synthesis were selected as representatives. These types include A/G, C/T, G/A, G/T, T/C mismatches and single-base deletion. Their mismatch recognition capabilities were quantitatively analyzed *via* electrophoretic mobility shift assay. As shown in **Fig. 1**, the binding rates of *Taq*MutS to A/G, C/T, G/A, G/T, T/C mismatches and single-base deletion were 78.6%, 36.8%, 59.1%, 59.9%, 69.7% and 71.7% respectively. These values were generally 10.2% to 17.4% lower than the corresponding binding rates of *Eco*MutS. Notably, the non-specific binding rate of *Taq*MutS to perfectly matched DNA was as high as 39.6%, which was significantly higher than that of *Eco*MutS (7.2%). Compared to *Eco*MutS, *Taq*MutS exhibited not only lower mismatch recognition efficiency but also stronger non-specific binding, which was consistent with previous findings.^9,15^ Nevertheless, *Taq*MutS possesses distinct advantages in biotechnological applications due to its superior thermostability. Therefore, we selected *Taq*MutS as the model to investigate the molecular mechanism of mismatch recognition.

**Fig. 1.**
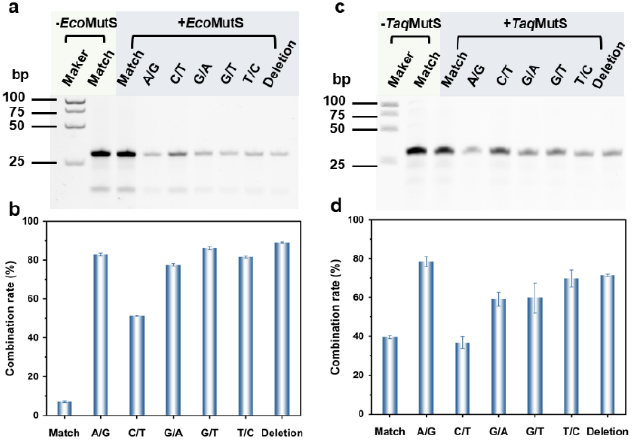
Analysis of mismatch recognition ability of *Eco*MutS and *Taq*MutS. (a,c) Non-denaturing polyacrylamide gel results and (b,d) gray-scale quantitative results of *Eco*MutS or *Taq*MutS binding to different mismatched DNAs.

Given that *Taq*MutS exhibited the highest binding affinity for A/G mismatches, this mismatch type was selected for computational simulation investigations. To identify the key residues involved in mismatched DNA recognition, we compared the dynamic conformational features and interaction energy differences between *Taq*MutS complexes with perfectly matched DNA and those with A/G-mismatched DNA. A triad region consisting of the core mismatched base pair (DG9:DT22) and its flanking adjacent base pairs (DC8:DG23 and DC10:DG21) was selected. Twenty-three residues within a 5 Å radius of this region were analyzed (**Fig. 2a**). Subsequently, for the complex systems of *Taq*MutS bound to perfectly matched DNA and mismatched DNA respectively, the molecular mechanics/generalized born surface area (MM/GBSA) free energy decomposition method was employed. Residues with |*ΔΔ*G| < 1 kcal/mol were excluded from further analysis (**Fig. 2b**). Additionally, combined with the evolutionary conservation analysis results of the MutS family (**Fig. 2c**), highly conserved functional sites (F39, Y40, and E41) were excluded. Finally, six potential sites that may affect the mismatch-binding ability of *Taq*MutS were identified, namely V36, V57, K61, R76, D472, and R473. Furthermore, alanine scanning mutagenesis was used to evaluate the impact of these sites on the mismatch recognition capability of *Taq*MutS. As shown in **Fig. S3** and **S4**, the binding rates of all mutants to perfectly matched DNA were significantly reduced (< 20%). This indicated that these residues are involved in the non-specific binding of *Taq*MutS. In addition, all mutants exhibited significant defects in mismatch-binding ability, particularly mutants V36A and V57A (**Fig. S4a** and **S4b**). Specifically, V36A completely lost its binding ability to A/G and C/T mismatches (binding rate < 10%). Its binding efficiency to other mismatches (G/T, T/C, and single-base deletion) was also reduced to 40-65%. Mutant V57A only retained binding ability to T/C mismatches (65.0%) and single-base deletions (64.2%), while its binding activity to other mismatches (A/G, C/T, G/T) was almost completely abolished. It should be noted that the D472A mutant exhibited severe aggregation after purification, which precluded the assessment of its mismatch recognition ability. Overall, these results demonstrated that V36, V57, K61, R76, and R473 are all key residues regulating the mismatch recognition function of *Taq*MutS.

**Fig. 2.**
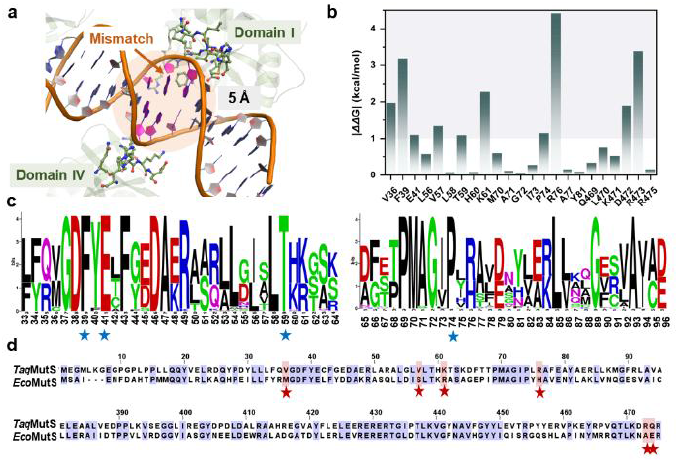
Alanine scanning mutagenesis and the design of homologous substitution mutation sites. (a) Structural visualization of amino acid residues within 5 Å of the mismatched base pairs and their adjacent base pairs. (b) Analysis of the free energy decomposition contribution of WT binding to correct and A/G mismatched DNA using the MM/GBSA binding method, where |*ΔΔ*G| = |*Δ*G_correct - *Δ*G_A/G mismatch|. (c) Sequence conservation analysis of the MutS family. (d) Comparative sequence analysis of *Taq*MutS and *Eco*MutS homologs.

Compared to *Taq*MutS, *Eco*MutS not only possesses higher binding affinity for various types of mismatched DNA but also shows a lower non-specific binding rate to perfectly matched DNA. Therefore, based on the homologous sequence alignment results between *Eco*MutS and *Taq*MutS (**Fig. 2d**), a homology substitution strategy was applied to the key residues to design six mutants, namely V36M, V57S, K61R, R76H, D472N, and R473A. The binding efficiency of these mutants to different types of DNA was then evaluated. As shown in **Fig. S4f-j**, the binding rate of V36M to perfectly matched DNA was reduced to 17.5%. However, its binding efficiency to C/T, G/T mismatches, and single-base deletion DNA was drastically decreased to less than 10% (**Fig. S4f**). For V57S, the non-specific binding efficiency was significantly reduced to 28.2%. Meanwhile, its recognition efficiency for A/G mismatch and single-base deletion DNA was notably enhanced, increasing to 98.4% and 94.9%, respectively (**Fig. S4g**). Although K61R maintained high binding efficiency (80-95.2%) for all types of mismatched DNA, its non-specific binding efficiency increased to 71.8% (**Fig. S4h**). Notably, D472N exhibited undifferentiated binding ability to all DNA types, with a binding efficiency as high as 98% for each. This result indicated that D472N completely lost the ability to distinguish between perfectly matched and mismatched DNA (**Fig. S4j**).

Based on the above findings, Gaussian accelerated molecular dynamics (GaMD) simulations were further performed to investigate the effects of mutations at these key sites on the conformational dynamics of *Taq*MutS. It has been reported that DNA mismatches can induce local conformational distortion of the double helix centered on the mismatched base pair. The mechanism by which *Taq*MutS recognizes mismatched DNA may be associated with this phenomenon.^19,20^ Therefore, focusing on the A/G mismatched base pair, we selected one adjacent base pair upstream and downstream of it. The number of hydrogen bonds and stacking angles formed among these three base pairs were analyzed to explore the local deformation of mismatched DNA in wild-type (WT)-*Taq*MutS and its mutants.

As shown in **Fig. S5** and **S6**, in the complex system of WT-*Taq*MutS with perfectly matched DNA, the number of hydrogen bonds among the three base pairs ranged from 7 to 9. Additionally, the stacking angles of bases DC8, DA9, and DC10 were distributed within the range of 150-180°, showing a parallel distribution trend. These results indicated that the three base pairs maintain a normal double-helix structure. In contrast, in the complex system of WT-*Taq*MutS with A/G-mismatched DNA, the number of hydrogen bonds among these three base pairs decreased to 4-7 (**Fig. 3a** and **3b**). For the V57S mutant, this number further reduced to 3-6 (**Fig. 3c** and **3d**), suggesting that the mismatched DNA undergoes local deformation at the mismatch site and the hydrogen bond interactions between mismatched base pairs are weakened. Furthermore, in the WT-*Taq*MutS/A/G-mismatched DNA complex, the π-stacking angles between the mismatched base DG9 and its adjacent upstream/downstream bases were mainly concentrated at 125° and 150° (**Fig. 3e** and **3f**). In the V57S mutant, however, the π-stacking angles were predominantly centered at 120° (**Fig. 3e** and **3g**). This observation demonstrated that the parallel stacking between bases was disrupted and the mismatched DNA exhibited local conformational distortion. Taken together, these results confirmed that the local bulging deformation of mismatched DNA was a key factor for *Taq*MutS recognition, and the degree of this deformation was closely correlated with the binding rate of *Taq*MutS to mismatched DNA.

**Fig. 3.**
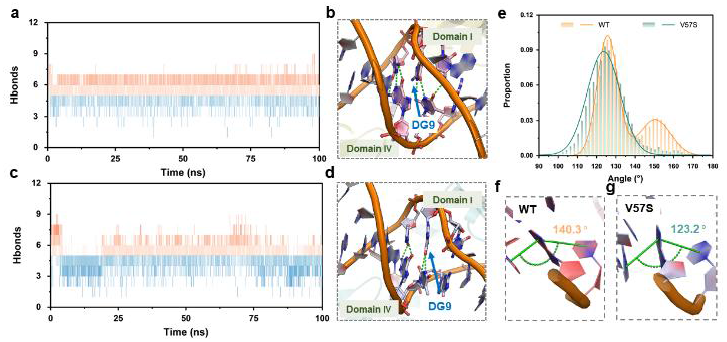
Local deformation analysis of mismatched DNA in the WT and V57S systems. (a) The number of hydrogen bonds between DC8, DA9, and DC10 and their paired bases in WT and A/G mismatched DNA systems during 100 ns GaMD simulation and (b) the visualized structure. (c) The number of hydrogen bonds between DC8, DA9, and DC10 and their paired bases in the V57S and A/G mismatched DNA systems during 100 ns GaMD simulation and (d) the visualized structure. (e) The probability distribution of π-stacking angles between the DC8, DA9, and DC10 base loops and the visualized structure of the (f) WT and the (g) V57S.

In the complex system of WT-*Taq*MutS and mismatched DNA, we observed that the distance between V57@CG and DG9@P was distributed in the range of 5-6 Å (**Fig. 4a** and **4b**). In contrast, in the V57S mutant system, the distance between S57@OG and DG9@P ranged from 2-5 Å. This phenomenon was attributed to the formation of hydrogen bond interactions between the side-chain hydroxyl group of S57 and the phosphate group on the backbone of mismatched DNA (**Fig. 4a** and **4c**). Considering that V/S57 is spatially adjacent to F39, the hydrogen bond interaction between S57 and mismatched DNA could drive the mismatched DNA to move closer to F39. Previous studies have confirmed that F39 is a key residue for mismatch recognition, prompting us to further analyze the interaction between F39 and mismatched DNA. In the V57S-mismatched DNA complex, the distance between the benzene ring of F39 and the base ring of DC8 was approximately 4 Å, while that between the benzene ring of F39 and the base ring of DG9 was maintained at around 6 Å (**Fig. 4d-f** and **S7**). However, in the WT complex, the distance between the benzene ring of F39 and the base rings of both DC8 and DG9 was maintained at approximately 7 Å (**Fig. 4d** and **4g**). These results indicated that the π-stacking interaction between F39 and the DNA mismatch site was stronger in V57S. This effect was ascribed to the hydrogen bond formed by S57 and mismatched DNA, which shortened the intermolecular distance and thus stabilized the binding of F39. Notably, in the WT-perfectly matched DNA complex, the distance between the benzene ring of F39 and the base of DA9 exceeded 10 Å (**Fig. S8**). This finding further demonstrated that the specific recognition of mismatched DNA by *Taq*MutS is closely associated with the interaction between F39 and the mismatch site.

**Fig. 4.**
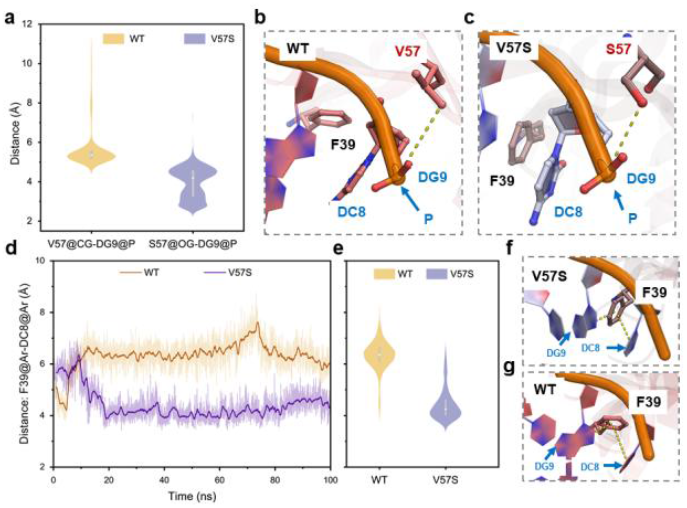
Interaction analysis of V/S57 and F39 with A/G mismatched DNA in WT and V57S systems. (a) The distance distribution between V/S57 and mis-matched base DG9 during 100 ns GaMD simulation, and visualized structure of (b) WT and (c) V57S. (d) The changes in distance and (e) the distance distribution between the benzene ring of F39 and the base ring of DC8 during 100 ns GaMD simulation, and visualized structural of (f) V57S and (g) WT.

Furthermore, D472N lost the ability to discriminate between perfectly matched DNA and mismatched DNA, with its non-specific binding efficiency exceeding 98%. This observation indicated that this residue plays a particularly critical role in mediating the non-specific recognition of *Taq*MutS. In the WT system, the distance between D472 and the perfectly matched DNA strand exceeded 8 Å (**Fig. S9a** and **S9b**). By contrast, in the D472N mutant, N472 formed novel hydrogen bond interactions with DG23 (the base paired with DC8), with the intermolecular distance maintained below 4 Å (**Fig. S9a** and **S9c**). Additionally, the contact frequency between D472 and mismatched DNA in the WT system was less than 20%, whereas the contact frequency between N472 and mismatched DNA in the D472N system exceeded 75%, leading to an even lower contact frequency between DNA and F39 (**Fig. S9d-f**). This conformational change impaired the ability of *Taq*MutS to distinguish between perfectly matched and mismatched DNA. To verify whether this effect depended on the negatively charged side chain, we constructed the N472E mutant by substituting the uncharged asparagine with negatively charged glutamic acid. As shown in **Fig. S4k**, N472E restored the capacity to recognize mismatched DNA, confirming that the DNA-repelling effect mediated by the negatively charged side chain is essential for the mismatch recognition function of *Taq*MutS. Collectively, these results demonstrated that the distance between residue 472 and mismatched DNA was inversely correlated with the distance between residue 57 and mismatched DNA (**Fig. 5a-c**). Specifically, for V57S, which exhibited superior mismatch recognition ability, the mismatched DNA was positioned closer to S57 and F39 but farther from D472 (**Fig. 5b**). For D472N, which failed to distinguish between perfectly matched and mismatched DNA, the mismatched DNA was located farther from S57 and F39 but closer to N472 (**Fig. 5c**). Thus, we proposed that S57 and D472 synergistically regulate the distance between F39 and mismatched DNA through a push-pull mechanism, thereby modulating the mismatch recognition capability of *Taq*MutS.

**Fig. 5.**
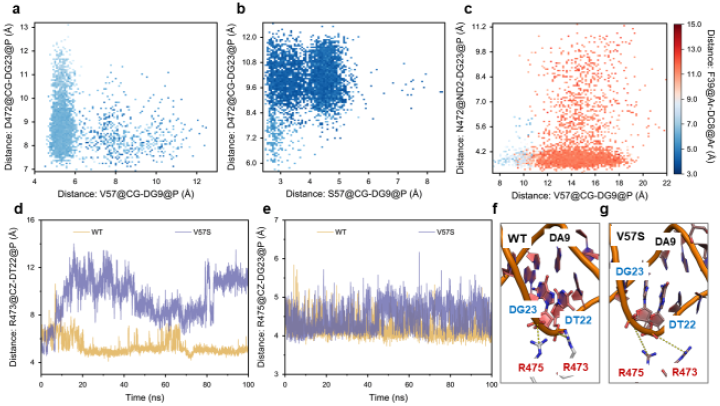
Analysis of the synergistic interaction of S57 and D472, and the mechanism of nonspecific binding of V57S. The distance distribution between V/S57 and DG9, D/N472 and DG23, as well as F39 and DC8 in the system of (a) WT, (b) V57S, and (c) D472N with A/G mismatched DNA during 100 ns GaMD simulations. The change in distance between (d) R473 and (e) R475 and the base loops of DT22 and DG23, respectively, in the system of WT and V57S with perfectly paired DNA, along with the visualized structures of (f) WT and (g) V57S.

To elucidate the molecular mechanism underlying the reduced non-specific binding of V57S, we constructed complex systems of WT and V57S bound to perfectly matched DNA, respectively, followed by 100 ns GaMD simulations. The results revealed that R473 and R475 of WT *Taq*MutS formed stable hydrogen bond interactions with perfectly matched DNA (**Fig. 5d-f**). The distance between R473@CZ and DT22@P ranged from 4 to 5 Å, and the distance between R475@CZ and DG23@P was maintained at 4 Å. In contrast, these two distances in V57S were distributed at 7-12 Å and 5 Å, respectively. This observation indicated the loss or attenuation of such hydrogen bond interactions, which contributed to the decreased binding rate of V57S to perfectly matched DNA (**Fig. 5d-g**). In addition, the binding rate of K61R to perfectly matched DNA increased from 39.6% to 71.8% (**Fig. S4h**), suggesting that this residue is involved in regulating the non-specific recognition of *Taq*MutS. Contact frequency analysis showed that in the WT system, K61 exhibited a relatively low contact frequency with the DNA base moiety but could form hydrogen bonds with phosphate groups to stabilize the DNA structure (**Fig. 6a-c**). On the contrary, in K61R, R61 displayed a higher contact frequency with DNA bases and no interaction with phosphate groups. This phenomenon might be attributed to the longer side chain of arginine, which hinders the formation of hydrogen bonds with DNA due to steric hindrance effects (**Fig. 6a-d**).

**Fig. 6.**
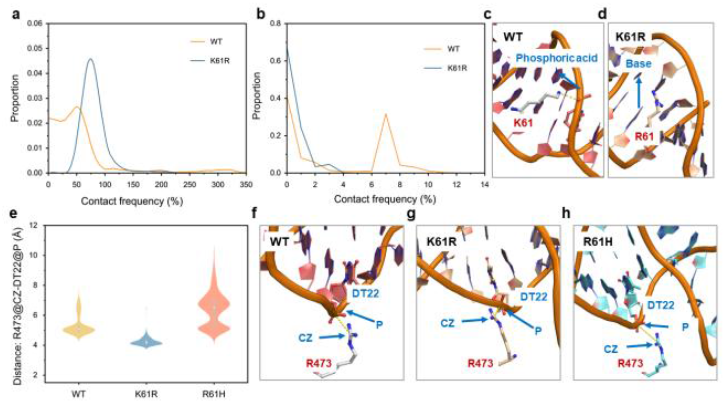
Analysis of the mechanism of electrostatic synergy between K61 and R473. (a) The frequency distributions of K/R61 contacts with the base portion and (b) the phosphate group portion of DNA, respectively, in the system of WT and K61R with perfectly paired DNA, and visualized structural of (c) WT and (d) K61R. (e) The distance distribution between R473 and DT22 during 100 ns GaMD simulation, and visualized structural of (f) WT, (g) K61R, and (h) R61H.

Given that K/R61 and R473 are located on opposite sides of the DNA strand (**Fig. S10**), they engage in competitive binding to the DNA phosphate backbone. In the WT system, the distance between R473@CZ and DT22@P was found to be above 5 Å, whereas this distance was maintained at approximately 4 Å in the K61R system (**Fig. 6e-g**). These results suggested that perfectly matched DNA, lacking electrostatic interactions with R61, instead forms hydrogen bond interactions with R473 on the opposite side, thereby enhancing the non-specific binding of K61R. To reduce R61’s steric hindrance, maintain its electrostatic interactions with DNA and keep DNA away from R473, we constructed the R61H mutant. In the R61H system, the distance between R473@CZ and DT22@P ranged from 5 to 8 Å (**Fig. 6e** and **6h**). This finding indicated the disappearance of hydrogen bonds between perfectly matched DNA and R473. This structural change corresponded to the reduction of R61H’s binding rate to perfectly matched DNA to below 20%, demonstrating that K61 is involved in the non-specific binding of *Taq*MutS (**Fig. S4l**).

This study clarified the DNA mismatch recognition mechanism of *Taq*MutS. Local conformational strain of mismatched DNA serves as the structural prerequisite. S57 enhances DNA binding through novel hydrogen bonds to stabilize F39-mismatch π-stacking interactions, whereas D472 ensures precise DNA positioning via electrostatic repulsion, thereby facilitating F39-mediated mismatch recognition. K61 and R473 dynamically regulate non-specific binding to perfectly matched DNA through electrostatic coordination. Taken together, these findings deepen the mechanistic understanding of *Taq*MutS function and provide a theoretical basis for the rational design of highly efficient and specific mismatch recognition proteins for biotechnological applications.

Wenyan Jiang: data curation, writing – original draft, software, formal analysis and investigation. Yiyang Ding: methodology, visualization and validation. Yuxuan Li: validation and visualization. Haojie Sun: visualization. Xiao Liang: conceptualization, funding acquisition, supervision and writing – review & editing. Quanshun Li: resources, funding acquisition, project administration and writing – review & editing.

This work was supported by the National Key R&D Program of China (2020YFA0907000), the National Natural Science Foundation of China (U24A20365, 32301051 and 32271319), the Science and Technology Department of Jilin Province (20250205039GH), the Development and Reform Commission of Jilin Province (2023C015), the China Postdoctoral Science Foundation (2025T180729) and the Fundamental Research Funds of the Central Universities, China (2024-JCXK-11).

## Conflicts of interest

There are no conflicts to declare.

## Data availability

The data supporting this article have been included as part of the Supplementary Information. Supplementary information is available. See DOI: [URL – formathttps://doi.org/DOI]

Notes and references

## Notes

### Competing Interest Statement

The authors have declared no competing interest.

