## Supplementary information for "Computer-assisted dissection of the DNA mismatch recognition mechanism of the MutS protein"

**Materials.**

Tryptone, yeast extract, *Escherichia coli* (*E. coli*) DH5α, *E. coli* BL21(DE3), single-stranded DNA (ssDNA) for mismatched DNA construction, and primers for site-directed mutagenesis were purchased from Sangon Biotech (Shanghai, China). Codon optimization and gene synthesis were provided by GenScript (Nanjing, China). DNA Polymerase was obtained from (SparkJade, Shandong, China). Restriction endonuclease DpnI was purchased from Thermo Fisher (Waltham, MA, USA). Isopropyl β-D-1-thiogalactopyranoside (IPTG) was purchased from Sigma-Aldrich (Shanghai, China). 10× DNA/RNA Native Sample Loading Buffer was obtained from Beyotime (Shanghai, China).

**Construction of strain and plasmid**

The DNA sequences of *Taq*MutS (GenBank accession No.: WP_053767724) and *Eco*MutS (GenBank accession No.: AAC75775) were synthesized by GenScript. These sequences were inserted into the pET-32a(+) plasmid, and the resulting recombinant plasmids were transformed into *E. coli* BL21(DE3) and DH5α, respectively. After transformation, the bacterial cells were screened on LB medium supplemented with ampicillin.

**Site-directed mutagenesis**

Mutants were constructed using a site-directed mutagenesis kit. Mutagenic primers were designed to target the desired mutation sites, and one-step PCR was carried out with the *Taq*MutS-encoding plasmid as the template to generate mutant plasmids. The primers are listed in **Table S1**. The main PCR procedure was as follows: pre-denaturation at 98 °C for 4 min; followed by 35 cycles of denaturation at 98 °C for 15 s, annealing at 75 °C for 15 s, and extension at 72 °C for 5 min. After PCR amplification, the products were digested with DpnI at 37 °C for 1 h, and then transformed into competent *E. coli* BL21(DE3) cells. After confirming the successful mutation by DNA sequencing, the mutant proteins were subjected to expression and purification.

**Molecular dynamics (MD) simulation**

The three-dimensional structures of the systems consisting of MutS proteins (including their variants) and mismatched DNA were constructed using AlphaFold3. Gaussian accelerated molecular dynamics (GaMD) simulations for all model systems were performed using the Graphics Processing Unit (GPU)-accelerated version of Amber 20. The ff14SB force field, OL15 force field, and General Amber Force Field (GAFF) were employed to generate the parameters and topological structures of proteins and DNA, respectively. A water box was added to each model using the TIP3P force field, with a minimum distance of 10 Å maintained between the enzyme surface and the edge of the water box. Sodium ions were also added to ensure the electrical neutrality of the system. Prior to formal simulations, all systems underwent energy minimization using the steepest descent method and conjugate gradient algorithm. After minimization, each system was gradually heated from 0 K to 298 K *via* Langevin dynamics, followed by equilibrium in the NVT (constant number of particles, volume, and temperature) ensemble. Additionally, the CPPTRAJ module in AmberTools20 and VMD software were used for the analysis and visualization of all MD simulation trajectories.

**Construction of mismatched DNA**

R-strand DNAs containing different types of mismatches and correct F-strand DNAs were separately dissolved in TE buffer (10 mM Tris, 0.1 mM EDTA, pH 7.5) to prepare single-stranded DNA (ssDNA) solutions with a concentration of 10 μM. Subsequently, the R-strand and F-strand ssDNA solutions were mixed at a volume ratio of 1:1. The mixture was treated at 95 °C for 5 min in a PCR thermocycler, then cooled down to 25 °C at a rate of 5 °C per minute and held at 25 °C for 5 min. Double-stranded DNA (dsDNA) solutions containing mismatches were obtained through this gradual annealing process. The sequences of R-strand and F-strand DNAs are listed in **Table S2**.

**Expression and purification of MutS protein**

*E. coli* cells harboring the MutS gene were inoculated into 50 mL of TB medium supplemented with 100 μg/mL ampicillin, and cultured at 37 °C and 180 rpm for 4 h until the $OD_{600}$ value reached 0.6. Subsequently, the culture temperature was adjusted to 18 °C, and 1 mM IPTG was added to induce protein expression, followed by continuous cultivation for 18 h. Bacterial cells were harvested by centrifugation at 8000× g for 10 min at 4 °C, and the pellet was resuspended in Tris-HCl buffer (20 mM, pH 7.5). Cells were lysed using an ultrasonic cell disruptor. After centrifugation of the lysate to remove impurities, the MutS protein with a 6×His tag was purified by affinity chromatography using Ni-NTA agarose resin. The purity of the purified protein was analyzed by sodium dodecyl sulfate-polyacrylamide gel electrophoresis (SDS-PAGE), and the protein concentration was determined by the Bradford method.

**Electrophoretic mobility shift assay**

Briefly, a reaction mixture was prepared by combining 10 μL of MutS protein solution (0.5 mg/mL), 2 μL of mismatched DNA solution, 2 μL of binding buffer (0.1 M KCl, 1 mM $MgCl_2$, 0.25 mM EDTA, 0.5 mM DTT, 5% v/v glycerol, 50 mM Tris-HCl, pH 8.5), and 8 μL of ultrapure water. The mixture was incubated at 37 °C for 30 min to ensure sufficient binding between MutS protein and dsDNA. After incubation, 10× DNA/RNA Native Sample Loading Buffer was added to the reaction system as recommended by the reagent manufacturer. The samples were loaded onto a 10% non-denaturing polyacrylamide gel (prepared with 0.5× TBE buffer) and electrophoresed at a constant voltage of 100 V for 2 h at 4 °C. Thereafter, the gel was stained by soaking in Gelblue nucleic acid dye (5× concentration) for 10 min, followed by decolorization with deionized water until the background became clear. A 200 bp DNA marker (Sangon Biotech, Shanghai, China) was used as the molecular weight reference. Gel images were captured using a Universal Hood II Gel Imaging System (Bio-Rad, USA). Gray-scale analysis of the gel images was performed using ImageJ 1.53 software. The grayscale value of the DNA band without MutS protein was set as 100%, and the binding rate of MutS protein to different dsDNA samples in each experimental group was calculated accordingly. All experiments were repeated three times, and the results were expressed as mean ± standard deviation.

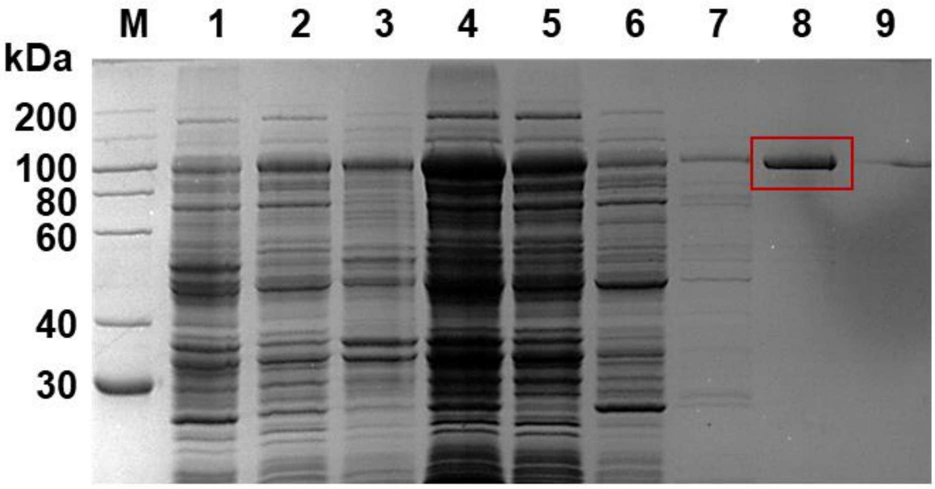


**Figure S1.** SDS-PAGE analysis of *Eco*MutS expressed in *E. coli* BL21(DE3). M: protein marker; 1: uninduced bacterial cell; line 2: IPTG-induced bacterial cell; 3: sediment of IPTG-induced bacterial cell lysate; 4: supernatants of IPTG-induced bacterial cell lysate; line 5: effluent fractions of the supernatant obtained from the lysate of IPTG-induced bacteria; 6: the fraction eluted with 10 mM imidazole; 7: the fraction eluted with 50 mM imidazole; 8: the fraction eluted with 150 mM imidazole; 9: the fraction eluted with 500 mM imidazole.

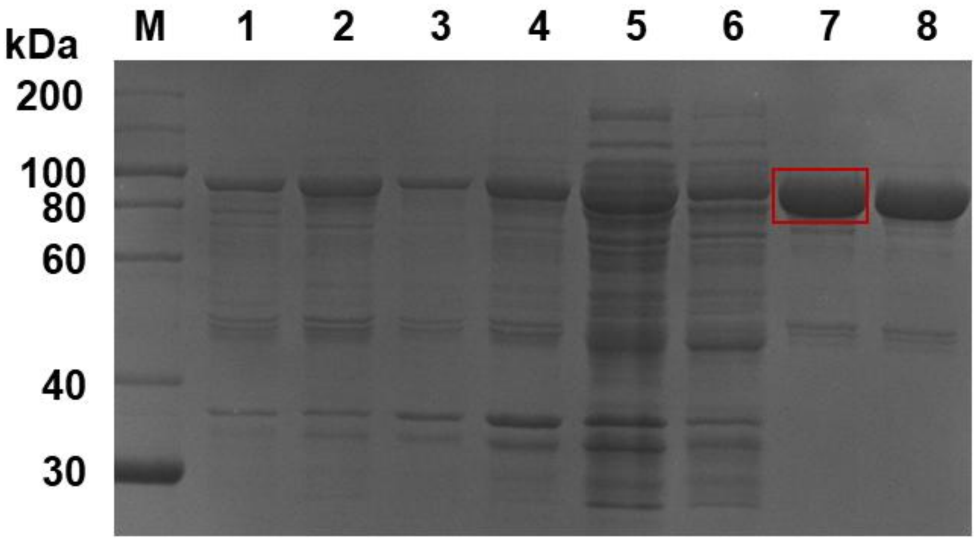


**Figure S2.** SDS-PAGE analysis of *Eco*MutS expressed in *E. coli* BL21(DE3). M: protein marker; 1: uninduced bacterial cell; line 2: IPTG-induced bacterial cell; 3: sediment of IPTG-induced bacterial cell lysate; 4: supernatants of IPTG-induced bacterial cell lysate; line 5: effluent fractions of the supernatant obtained from the lysate of IPTG-induced bacteria; 6: the fraction eluted with 50 mM imidazole; 7: the fraction eluted with 150 mM imidazole; 8: the fraction eluted with 500 mM imidazole.

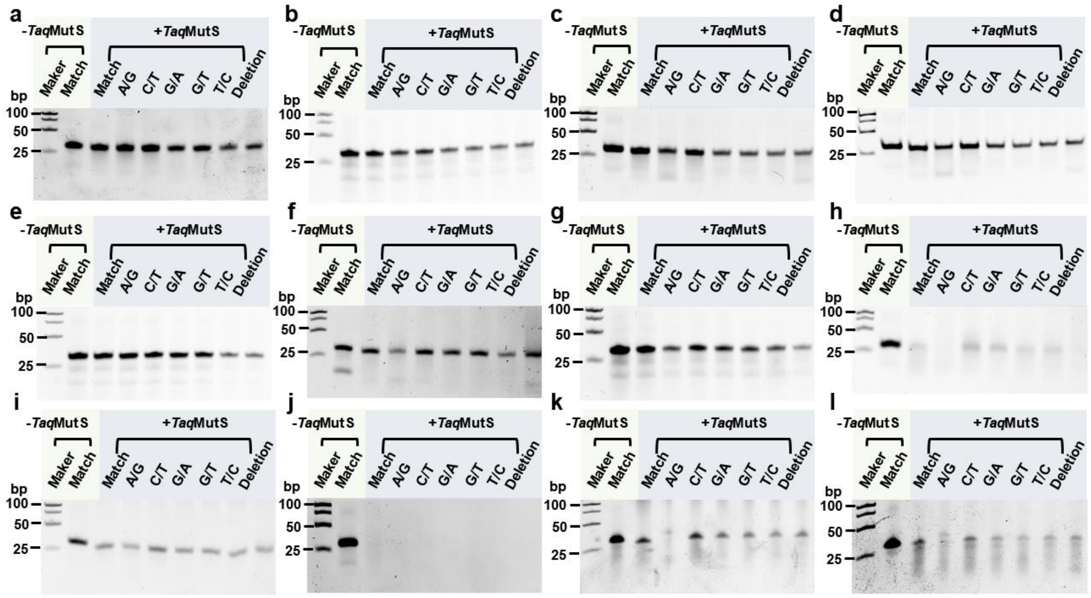

**Figure S3.** Non-denaturing polyacrylamide electrophoresis analysis of *Taq*MutS mutants for their binding ability to different mismatched DNAs. (a) V36A; (b) V57A; (c) K61A; (d) R76A; (e) R473A; (f) V36M; (g) V57S; (h) K61R; (i) R76H; (j) D472N; (k) N472E; (l) R61H.

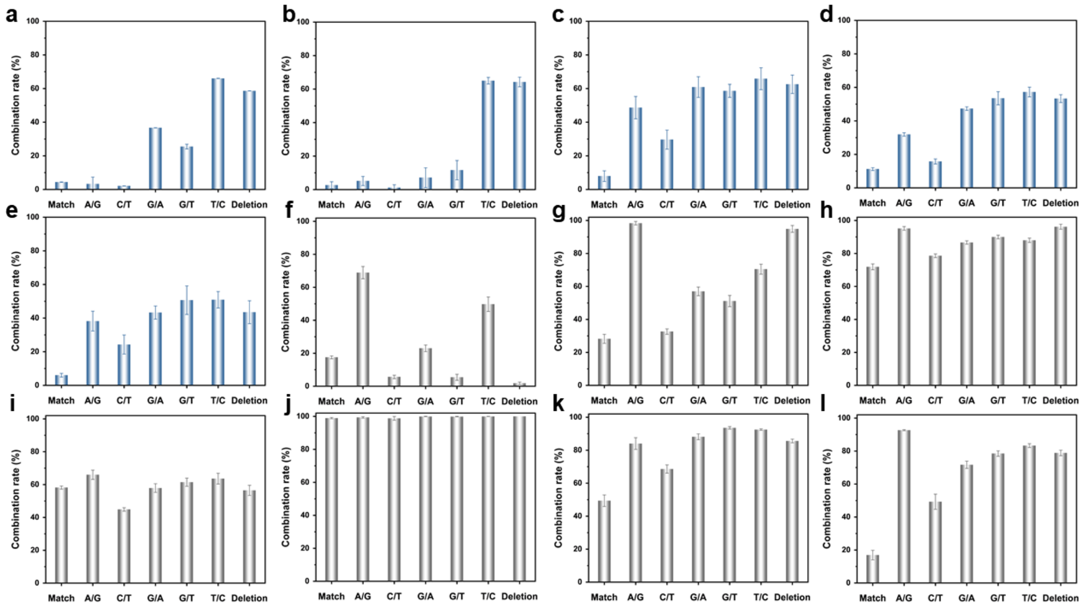


**Figure S4.** Gray-scale quantitative results of *Taq*MutS mutants binding to different mismatched DNAs. (a) V36A; (b) V57A; (c) K61A; (d) R76A; (e) R473A; (f) V36M; (g) V57S; (h) K61R; (i) R76H; (j) D472N; (k) N472E; (l) R61H.

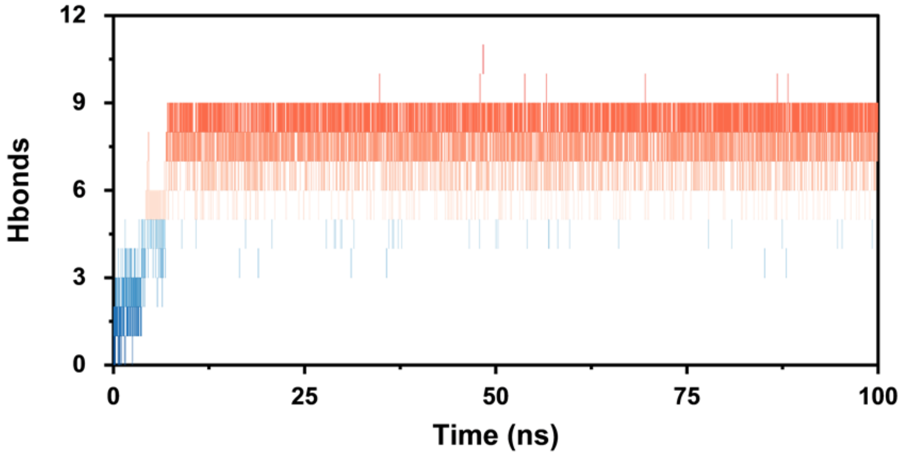


**Figure S5.** Hydrogen bond counts between DC8, DA9, DC10, and their paired bases during 100 ns GaMD simulations of WT and the correctly paired DNA system.

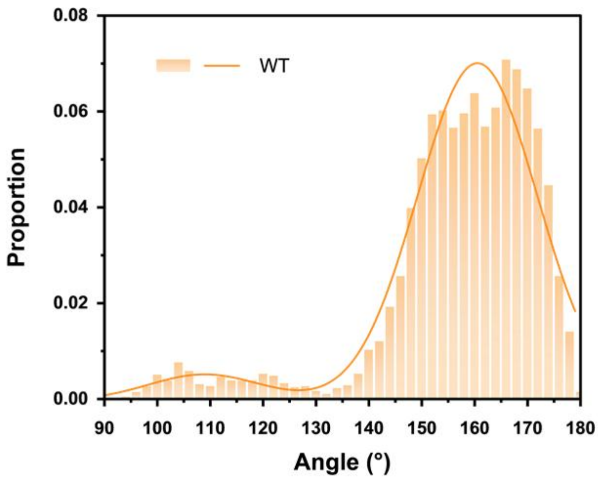


**Figure S6.** Probability distribution of π-stacking angles between WT and the DC8, DA9, and DC10 base loops within the correctly paired DNA system during 100 ns GaMD simulations.

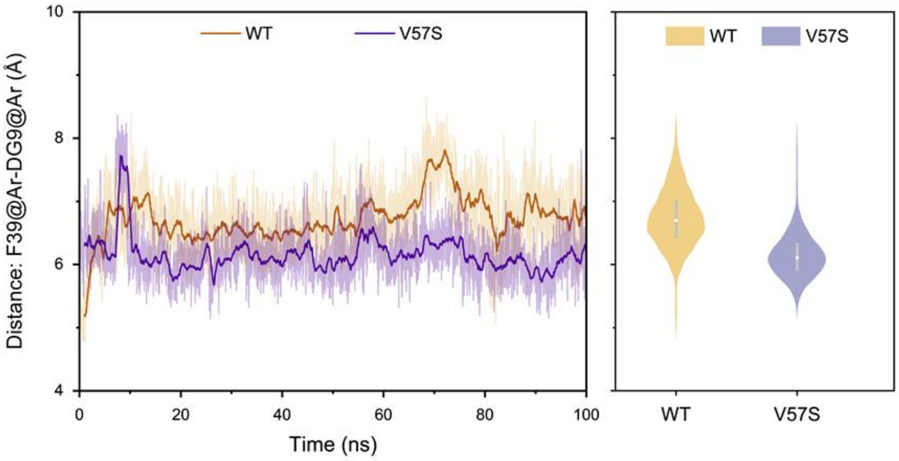


**Figure S7.** The changes in distance and the distance distribution between the F39 benzene ring and the DG9 base ring within the WT, V57S and mismatched DNA systems during 100 ns GaMD simulations.

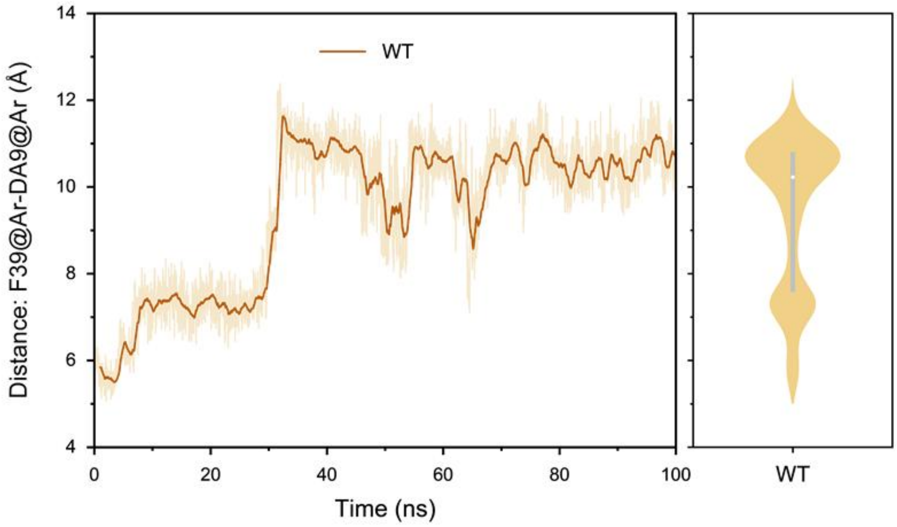


**Figure S8.** The changes in distance and the distance distribution between the F39 benzene ring and the DA9 base ring within the WT and correctly paired DNA systems during 100 ns GaMD simulations.

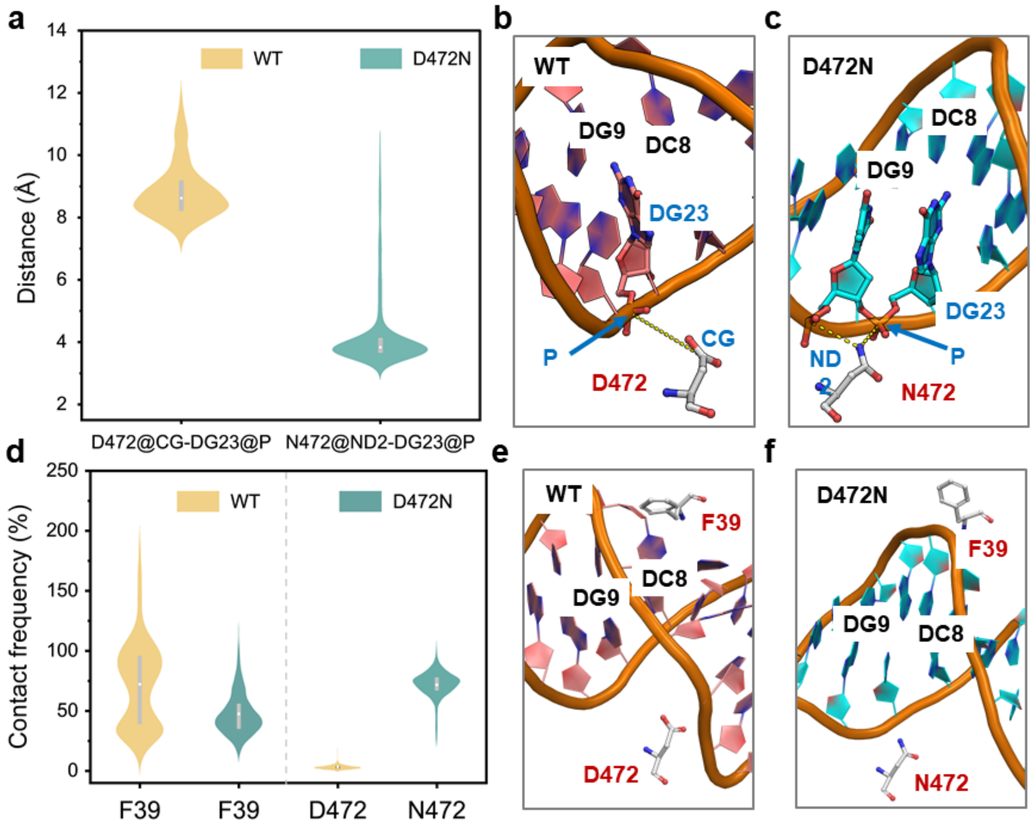


**Figure S9.** Interaction analysis of D/N472 and F39 with correctly matched DNA. (a) The distance distribution between D/N472 and base DG23 during 100 ns GaMD simulation, and the visualized structure of (b) WT and (c) D472N. (d) The contact frequencies of F39 and D/N472 with the correctly paired DNA during 100 ns GaMD simulation and the visualized structure of (e) WT and (f) D472N.

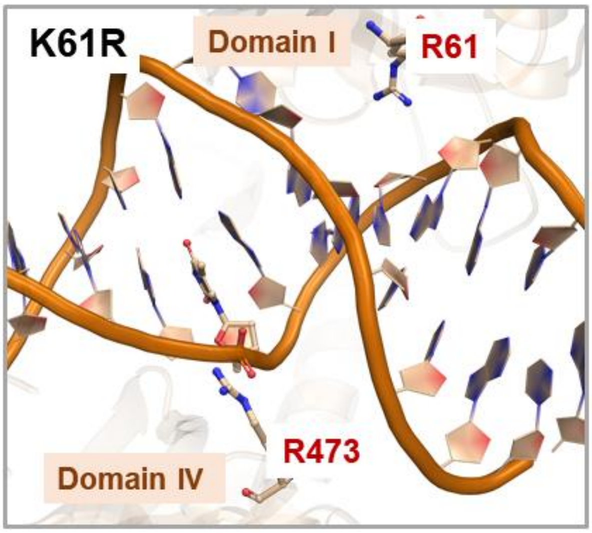


**Figure S10.** Schematic representation of the spatial distribution of R61 and R473 in K61R mutant.

**Table S1.** Primer sequences for mutant construction.

| Mutants | Primers |
|---|---|
| V36A | CTTGTTGTTCCAGGCCGGTGATTTCTACGAG |
| | CTCGTAGAAATCACCGGCCTGGAACAACAAG |
| V36M | CTATCTCTTGTTGTTCCAGATGGGTGATTTCTACGAGTGC |
| | GCACTCGTAGAAATCACCCATCTGGAACAACAAGAGATAG |
| V57A | GCGCTAGGACTGGCGCTGACCCATAAAAC |
| | GTTTTATGGGTCAGCGCCAGTCCTAGCGC |
| V57S | GTGCGCTAGGACTGAGCCTGACCCATAAAAC |
| | GTTTTATGGGTCAGGCTCAGTCCTAGCGCAC |
| K61A | CTGGTCCTGACCCATGCGACCAGCAAAGACTTC |
| | GAAGTCTTTGCTGGTCGCATGGGTCAGGACCAG |
| K61R | GACTGGTCCTGACCCATCGTACCAGCAAAGACTTCAC |
| | GTGAAGTCTTTGCTGGTACGATGGGTCAGGACCAGTC |
| R61H | GACTGGTCCTGACCCATCATACCAGCAAAGACTTCAC |
| | GTGAAGTCTTTGCTGGTATGATGGGTCAGGACCAGTC |
| R76A | CGGGTATCCCGTTGGCGGCGTTTGAGGCGTAC |
| | GTACGCCTCAAACGCCGCCAACGGGATACCCG |
| R76H | GGTATCCCGTTGCATGCGTTTGAGGC |
| | GCCTCAAACGCATGCAACGGGATACC |
| D472A | GTTCAGACCTTGAAAGCGCGTCAGCGTTACACC |
| | GGTGTAACGCTGACGCGCTTTCAAGGTCTGAAC |
| D472N | CCAGTTCAGACCTTGAAAAATCGTCAGCGTTACACCCTTC |
| | GAAGGGTGTAACGCTGACGATTTTTCAAGGTCTGAACTGG |
| N472E | CAGACCTTGAAAGAACGTCAGCGTTACACC |
| | GGTGTAACGCTGACGTTCTTTCAAGGTCTG |
| R473A | CAGACCTTGAAAGACGCGCAGCGTTACACCCTTC |
| | GAAGGGTGTAACGCTGCGCGTCTTTCAAGGTCTG |

**Table S2.** The sequence of mismatched DNA.

| DNA | Sequences |
|---|---|
| Match-F | TATCCAGCTGCCAGTCACCAGTGTCAGCGT |
| Match-R | ACGCTGACACTGGTGACTGGCAGCTGGATA |
| C/T-R | ACGCTGACACTGGTGATTGGCAGCTGGATA |
| G/T-R | ACGCTGACACTGGTGACTTGCAGCTGGATA |
| G/C-R | ACGCTGACACTGGTGACTCGCAGCTGGATA |
| A/G-R | ACGCTGACGCTGGTGACTGGCAGCTGGATA |
| T/C-R | ACGCTGACACTGGTGACCGGCAGCTGGATA |
| Deletion-R | ACGCTGACACTGGTGA_TGGCAGCTGGATA |
